# Mitochondria affect photosynthesis through altered tissue levels of oxygen

**DOI:** 10.1101/2024.07.19.604342

**Authors:** Matleena Punkkinen, Olga Blokhina, Lucas León Peralta Ogorek, Minsoo Kim, Kurt Fagerstedt, Elizabeth Vierling, Ole Pedersen, Alexey Shapiguzov

## Abstract

Interactions between plant energy organelles, the chloroplasts and the mitochondria, are crucial for plant development and acclimation. These interactions occur at different levels including exchange of metabolites and reducing power, organelle signaling pathways and intracellular gas exchange. Mitochondrial retrograde stress signaling activates expression of nuclear genes encoding mitochondrial components, including alternative oxidases. High abundances of these respiratory enzymes coincide not only with the changes in plant respiration but also with alterations in the chloroplast. For example, plants that overexpress alternative oxidases are tolerant to methyl viologen, a redox-active compound that catalyzes transfer of electrons from Photosystem I to molecular oxygen. The mechanism of this inter-organelle interaction is unclear but could be related to diminished availability of tissue oxygen. Here we assessed respiration, photosynthesis and *in vivo* levels of oxygen in a set of Arabidopsis lines with perturbations in diverse mitochondrial functions, including defects in respiratory complex I, mitochondrial protein processing, transcription, nucleoid organization, altered fission and architecture or suppressed ATP synthase activity. In these lines, the increased abundance and activity of alternative oxidases strongly correlated with higher oxygen consumption in darkness, lower oxygen re-accumulation in light, and diminished effects of methyl viologen in chloroplasts. These results support the hypothesis that increased mitochondrial oxygen sink capacity affects photosynthesis by decreasing oxygen levels in tissues. This phenomenon can be one of the reasons for the impact that stressed mitochondria have on chloroplasts and photosynthesis. It contributes to our understanding of the mechanisms of hypoxia establishment and acclimation in plants.

## Introduction

Crosstalk between plant energy organelles, the chloroplasts and the mitochondria, is crucial for plant development and acclimation. The interactions between the organelles include exchange of metabolites and reducing equivalents (Yoshida and Noguchi, 2011; Vanlerberghe, 2013; Shameer et al., 2019; Møller et al., 2021) and retrograde signaling (de Souza et al., 2017; Waszczak et al., 2018; Shapiguzov et al., 2019; Wang et al., 2020). Another well-established interaction is the exchange of gases essential for photosynthesis and respiration. In chloroplasts, carbon dioxide (CO_2_) is converted to photoassimilates in the Calvin-Benson-Bassham cycle and molecular oxygen (O_2_) is released as a byproduct of the water splitting activity of the oxygen-evolving complex. O_2_ is consumed by photorespiration (Bauwe et al., 2010), activity of plastid terminal oxidase (Nawrocki et al., 2015) and the water-water cycle that is associated with production of reactive oxygen species (ROS) (Miyake, 2010). In mitochondria, CO_2_ is released as a byproduct of the tricarboxylic acid (TCA) cycle and its associated reactions and by photorespiratory glycine decarboxylation (Møller et al., 2021), while O_2_ is consumed as the final electron acceptor in the respiratory electron transfer chain by mitochondrial cytochrome *c* oxidase (COX) and alternative oxidase (AOX) pathways. AOXs catalyze oxidation of ubiquinol and reduction of O_2_ to water thereby acting as electron sinks (Vanlerberghe, 2013; Dahal and Vanlerberghe, 2017; Yoshida and Noguchi, 2011; Møller et al., 2021) and oxygen sinks (Gupta et al., 2009; Rasmusson et al., 2009; Kelliher and Walbot, 2012; van Dongen and Licausi, 2015). Intracellular and tissue gas exchange of O_2_ and CO_2_ has been thoroughly covered in the literature (Gerbaud and André, 1980; Peltier and Thibault, 1985; Parkhurst et al., 1988; Vesala et al., 1996; Hunt, 2003; Driever, 2024). However, molecular-level understanding of the implications of O_2_ distribution and fluxes within plant cells and tissues is still largely missing.

A ROS-dependent retrograde signaling pathway initiated in mitochondrial electron transfer chain induces expression of a set of nuclear genes called the mitochondrial dysfunction stimulon (MDS) that encodes AOXs and other, mostly mitochondrial, components (De Clercq et al., 2013; Ng et al., 2013; Shapiguzov et al., 2019; Khan et al., 2024). Activation of MDS and concomitant overaccumulation of AOXs not only changes plant respiration, but also modifies chloroplast metabolism and photosynthesis (Shapiguzov et al., 2019, 2020). A prominent feature of plants with activated MDS is their increased tolerance to methyl viologen (MV), also known as paraquat (De Clercq et al., 2013; Ng et al., 2013; Shapiguzov et al., 2019). MV catalyzes the transfer of electrons from Photosystem I (PSI) to O_2_, producing ROS (Miyake, 2010; Fitzpatrick et al., 2022). This is called the Mehler reaction, and it is the starting point of the water-water cycle. In physiological conditions it plays important roles in chloroplast signaling and metabolism. ROS produced in Mehler reaction oxidize chloroplast thiol redox enzymes, thereby controlling multiple thiol-redox pathways (Ojeda et al., 2018; Vaseghi et al., 2018; Yoshida et al., 2018). When the Mehler reaction is enhanced, e.g., with MV, the resulting generation of ROS can exceed the ROS-scavenging capacity of the chloroplast leading to inhibition of photosynthesis and eventually cell death (Farrington et al., 1973; Nishiyama et al., 2011; Hawkes, 2014; Shapiguzov et al., 2019).

Arabidopsis mutant *rcd1*, which has a constitutively active MDS response, is characterized by higher tolerance to MV and more reduced redox states of chloroplast thiol enzymes 2-Cys Peroxiredoxin and NTRC (Shapiguzov et al., 2019, 2020). Together, these phenotypes suggest that MDS overexpression affects the Mehler reaction in the chloroplast. The mechanisms of this inter-organelle interaction remain unclear. They have been linked to electron sink activity of mitochondrial AOXs, as these enzymes can consume reducing power transferred from chloroplasts to mitochondria via redox shuttles (Vanlerberghe, 2020; Shapiguzov et al., 2019). However, electron sink activities of AOXs cannot fully explain the interaction in darkness and under an hypoxic atmosphere (Shapiguzov et al., 2020). We recently proposed that this inter-organelle interaction is related to reduced availability of tissue O_2_ in plants with increased mitochondrial respiration (Shapiguzov et al., 2020). However, no evidence of lower O_2_ levels in mitochondrial mutants was available. Furthermore, RCD1 acts in numerous signaling pathways (Jaspers et al., 2009; Brosché et al., 2014; Shapiguzov et al., 2019), hence pleiotropic effects of the *rcd1* mutation on plant physiology prevented us from establishing more direct links between MDS overexpression and tissue O_2_ levels, changes in chloroplasts, and photosynthesis.

In this study we explored respiration, photosynthesis and *in vivo* O_2_ consumption / re-accumulation dynamics in a set of Arabidopsis mutants and transgenic lines with perturbations in mitochondrial functions (Fig. 1). These perturbations included reduced function of respiratory complex I – *ndufa1* (Meyer et al., 2011), *ndufs4* (Meyer et al., 2009) and *rug3* (Kühn et al., 2011); altered mitochondrial protein processing – *phb3* (Van Aken et al., 2007) and *lon1* (Rigas et al., 2009); deficiencies in mitochondrial transcription and nucleoid organization – *rpoTmp* (Kühn et al., 2009), *shot1* (Kim et al., 2012) and *atad3a1 atad3b1*:*ATAD3A1-GFP* (here called *ATAD3A1-GFP*; Kim et al., 2021); altered mitochondrial fission and architecture – *drp3a drp3b* (Fujimoto et al., 2009) and *mic60* (Michaud et al., 2016); or suppressed mitochondrial ATP synthase activity – *ATPd* RNAi line (here called *ATPQi*; Liu et al., 2021). We also included lines with altered mitochondrial retrograde MDS signaling: *rcd1*, *rcd1 anac017* (Shapiguzov et al., 2019), *ANAC013* overexpressor line (here called *ANAC013*-OE; De Clercq et al., 2013) and an *AOX1a* overexpressor line (here called *AOX1a*-OE; Umbach et al., 2005). AOX respiration affects mitochondrial production of nitric oxide (NO) (Vanlerberghe, 2013; Gupta et al., 2018). To address potential implication of NO in organelle interaction we included the *hot5* loss-of-function mutant deficient in S-nitrosoglutathione reductase (GSNOR), the enzyme affecting cellular NO levels, and a *GSNOR* overexpressor line (here called *GSNOR-GFP*; Feechan et al., 2005; Lee at al, 2008).

**Figure 1.**
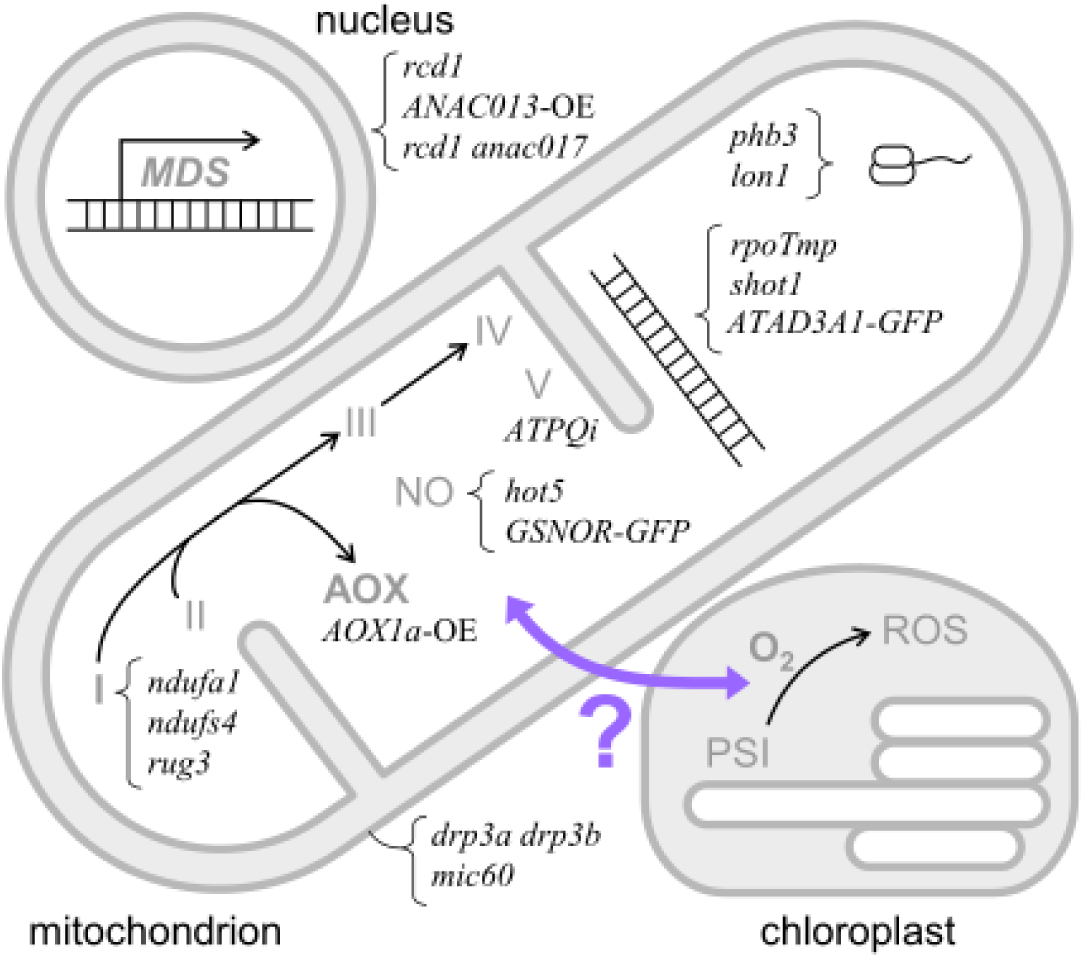
Diagram of mutants and transgenic lines used in the study. We utilized a set of mutants with a wide variety of mitochondrial perturbations to examine the interactions between mitochondrial respiration and chloroplastic Mehler reaction (purple arrow). The mutants and transgenic lines are marked in italics next to the affected functions, pathways or complexes.

Our results support the hypothesis that increased O_2_ sink activities associated with stressed mitochondria and accumulation of mitochondrial AOXs affect photosynthesis through decreasing tissue levels of O_2_. This phenomenon can be one of the reasons for the impact of stressed mitochondria on chloroplasts and photosynthesis. Our findings open new research possibilities to study hypoxia in plants. They will contribute to understanding the mechanisms of hypoxia establishment and sensing; plant signaling and adaptation to mitochondrial stress and hypoxia in various environmental and developmental contexts.

## Results

### Diverse types of mitochondrial defects increase abundance and respiration capacity of AOXs

Increased AOX abundance and enzymatic capacity coincide with changes in the chloroplasts of mutants with a constitutive MDS response (De Clercq et al., 2013; Ng et al., 2013; Shapiguzov et al., 2019, 2020). To find out whether this is a general phenomenon associated with various mitochondrial defects, we studied a large set of Arabidopsis mutants and transgenic lines with impaired mitochondrial functions (Fig. 1). We first evaluated abundance of AOX protein by immunoblotting protein extracts with an antibody recognizing all five isoforms of Arabidopsis AOX (Supplementary Figure 1A). Several of the tested lines demonstrated statistically significant increase in AOX abundance compared to the wild type (Fig. 2A). We next assessed AOX respiration capacity by measuring O_2_ consumption rate in darkness in the presence of KCN, the inhibitor of COX respiration.

**Figure 2.**
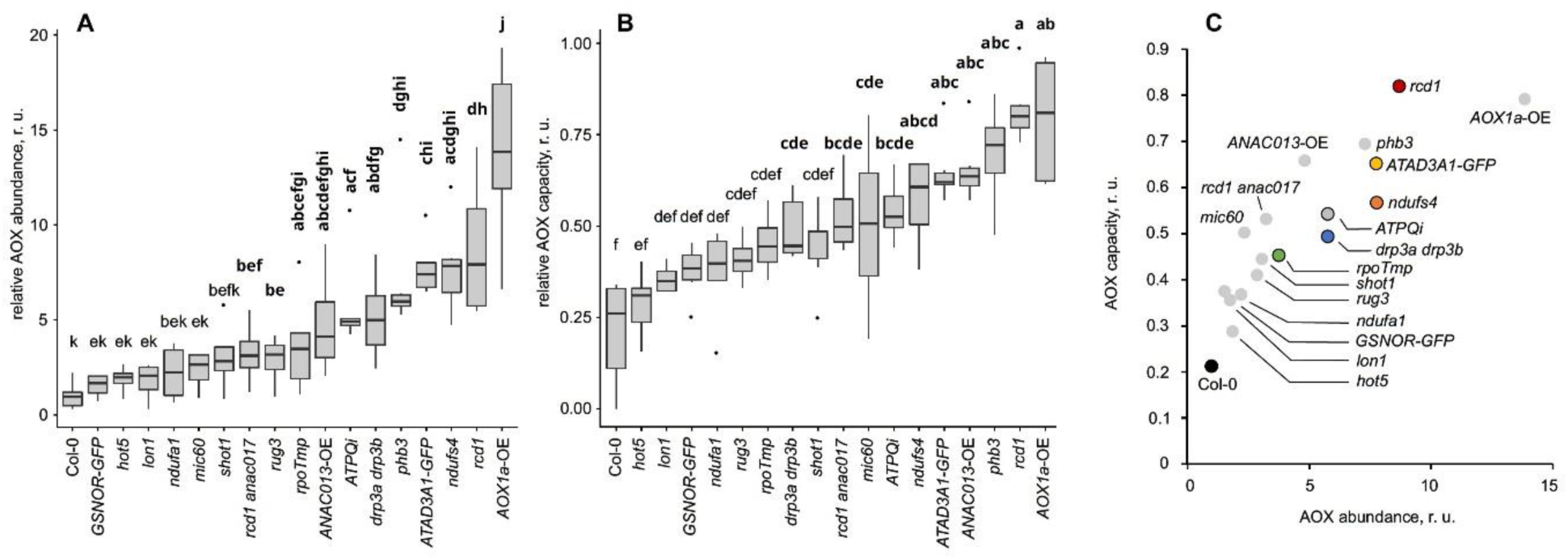
AOX abundance and capacity in plant lines with perturbed mitochondrial functions. AOX abundance was measured by immunoblotting total protein extracts from seedlings grown on plates (A), and AOX capacity was determined as the fraction of O_2_ consumption that was not inhibited by KCN (B). The genotypes are arranged in order of increasing mean value. Quantification in (A) was done using 4 independent immunoblots, and in (B) with 6 independent biological replicates. Statistically significant groups are indicated with letters, which are bolded for lines that have statistically significant difference from wild type. There was a strong correlation between AOX abundance and capacity (C), with Pearson correlation value of 0.8704 and p-value < 0.0001. The plant lines selected for in-depth analyses (see below) are marked with separately colored dots.

AOX activities are resistant to KCN, hence residual O_2_ uptake in KCN-treated seedlings can be ascribed to AOXs. In several lines, application of KCN led to only minor drops in O_2_ consumption, indicating high involvement of AOX respiration in the overall O_2_ uptake by the seedlings (Fig. 2B and Supplementary Figure 1B). Wild type plants (Col-0) had low AOX abundance and relatively low resistance to KCN, whereas the *AOX1a* overexpressor line and *rcd1* displayed both high AOX abundance and high KCN-resistant respiration. For all tested lines we found a strong correlation between AOX abundance and AOX capacity (Pearson correlation from averages 0.8704, p-value < 0.0001) (Fig. 2C).

### Mitochondrial perturbations modify plants’ responses to MV

To examine further the link between impaired mitochondrial function and MV resistance, we studied our plant lines’ tolerance to MV-induced photoinhibition. Leaf discs were floated overnight in darkness on solutions with or without MV to facilitate uptake of the chemical. After incubation, MV toxicity was induced by repetitive 1-hour light periods (450 nm, 80 µmol m^−2^ s^−1^) each followed by a 20 min dark period, then Fo and Fm measurement. The resulting decay of maximal quantum yield of Photosystem II (PSII) (Fv/Fm) was measured as described in (Shapiguzov et al., 2019). Importantly, after the overnight incubation and before the start of the light exposure the starting Fv/Fm was highly similar in MV-treated and control plants, in the range of 0.7-0.75 (Supplementary Figure 2). This indicated that the assessed effects of MV were light-dependent and thus likely caused by MV action in the chloroplast. Several lines, affected in different mitochondrial processes, were more tolerant to MV in comparison to the wild type (Fig. 3A, B). There was a strong correlation between tolerance to MV-induced photoinhibition and AOX abundance in the tested lines with one exception, the *AOX1a*-OE line (Fig. 3C).

**Figure 3.**
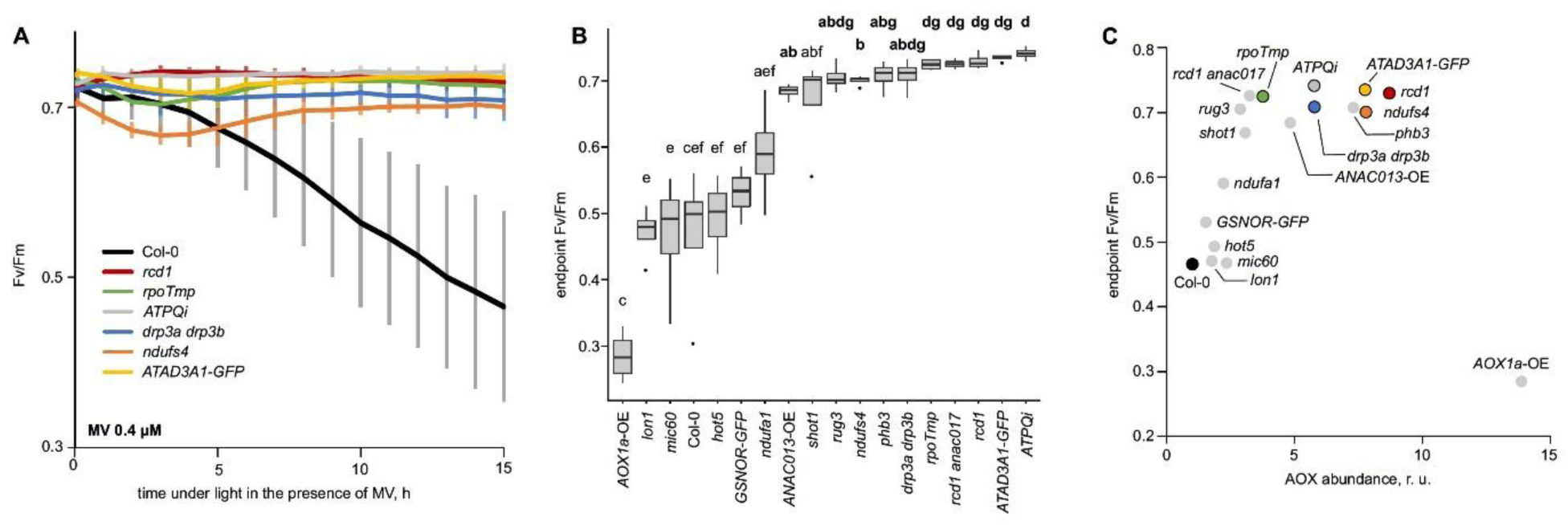
Responses of the plant lines with perturbed mitochondrial functions to MV-induced photoinhibition. Long-term exposure of MV-treated leaf discs to light led to gradual loss of maximal PSII quantum yield (Fv/Fm) in the Col-0 wild type, whereas MV-tolerant lines retained PSII function. The Fv/Fm dynamics of selected lines during long-term exposure to light is shown in (A). Shown values are averages +/-standard deviations. Quantification of Fv/Fm in all tested plant lines at the 15 h time point is shown in (B), with genotypes arranged in order of increasing mean. For each experiment we used leaf discs from at least 4 independent plants. The experiment was repeated twice with similar results. Statistically significant groups are indicated with letters, which are bolded for lines that have higher Fv/Fm than wild type. There was a strong correlation between tolerance to MV-induced photoinhibition and AOX abundance (C), with Pearson correlation value of 0.7478 and p-value < 0.001. The exception was the *AOX1a*-OE line, which was excluded from the calculation. The plant lines selected for in-depth analyses (see below) are marked with separately colored dots.

In addition to the long-term effects on chloroplasts and the photosynthetic apparatus associated with MV-induced ROS production, MV exerts rapid changes in the redox state of the photosynthetic electron transfer chain upon brief exposure to light. This effect of MV can be observed already after several hundred milliseconds of light exposure (Shapiguzov et al., 2020), and is likely explained by the fact that MV enhances the electron sink at the acceptor side of PSI, promoting oxidation of the electron transfer chain and thus faster quenching of chlorophyll fluorescence. We previously showed that in an aerobic atmosphere (normal air) the effect of MV on chlorophyll fluorescence quenching was indistinguishable between the wild type and *rcd1*, but in an hypoxic environment chlorophyll fluorescence quenching activity of MV diminished in *rcd1* while remaining high in the wild type (Shapiguzov et al., 2020). We therefore tested the effects of MV under an hypoxic atmosphere. First, we floated leaf discs overnight on solutions with or without MV in darkness under ambient atmosphere. Next, we introduced an hypoxic environment by flushing nitrogen gas over the leaf discs for 20 min in darkness. After this we performed light treatments and recorded the chlorophyll fluorescence response.

Firstly, we measured dark relaxation of maximal chlorophyll fluorescence (Fm) triggered by a saturating light pulse (Fig. 4A, C). In the second part of the assay we followed dynamics of chlorophyll fluorescence under low-intensity actinic light (Fig. 4B, D). In both types of measurements, chlorophyll fluorescence responses were similar in all the lines in aerobic environment (Supplementary Figure 3A, B) and in hypoxic atmosphere without MV (Supplementary Figure 3C). However, in hypoxic conditions the effect of MV on quenching was strikingly diverse between the genotypes (Fig. 4). In the wild type plants, MV still efficiently quenched chlorophyll fluorescence, while in several lines with impaired mitochondria the fluorescence dynamics of MV-treated samples became indistinguishable from MV-untreated controls (compare Fig. 4B to Supplementary Figure 3C). Interestingly, in these assays performance of the *AOX1a*-OE line was similar to that of the wild type, suggesting that enhanced expression of a single AOX1a isoform was not sufficient for the studied organellar interaction.

**Figure 4.**
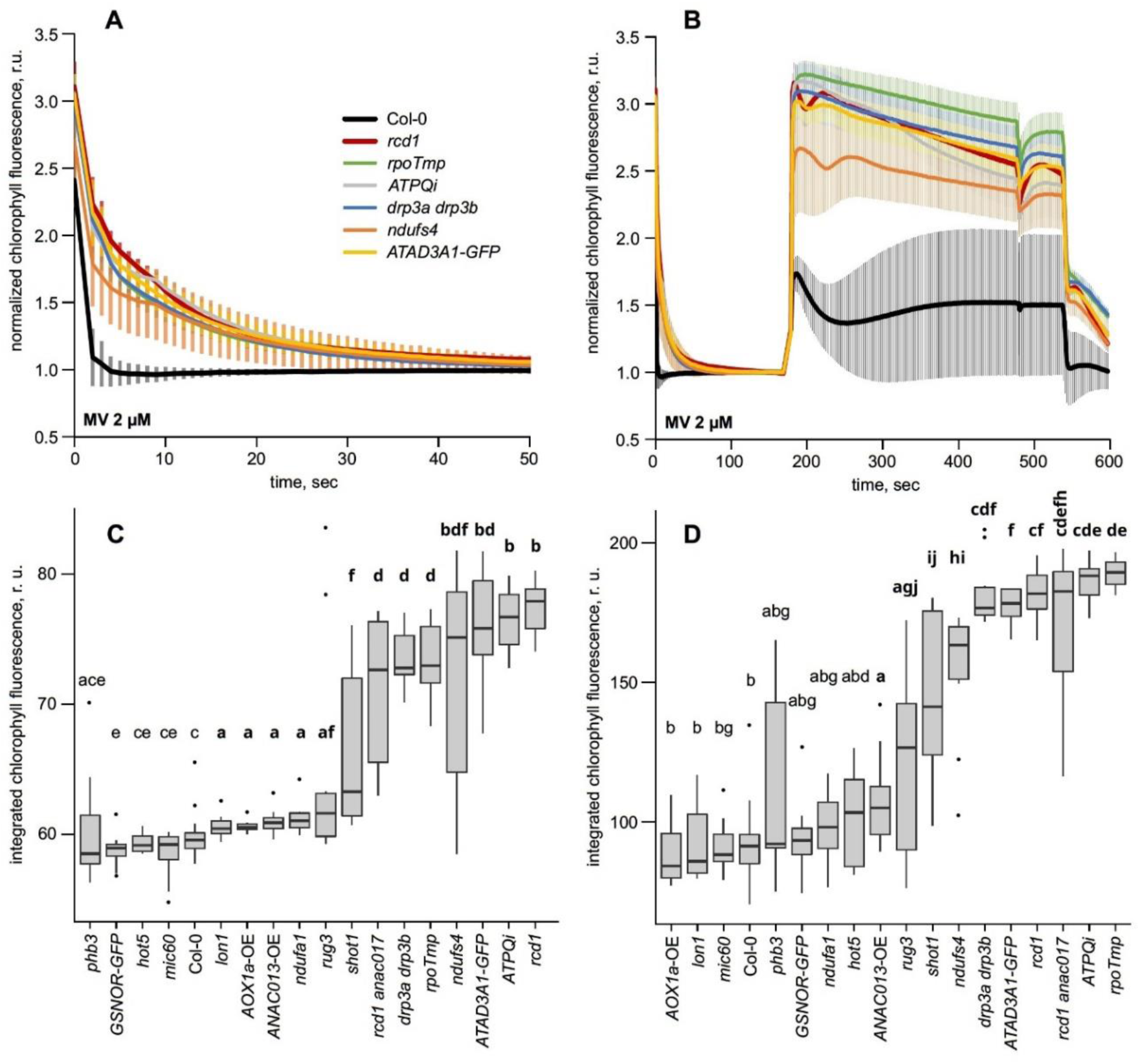
Responses of plant lines with perturbed mitochondrial functions to MV-induced quenching of chlorophyll fluorescence under hypoxia. In comparison to MV-sensitive lines such as Col-0 wild type, several plant lines displayed resistance to MV-induced chlorophyll fluorescence decay under an hypoxic atmosphere. This effect was observable in both dark relaxation of Fm (A, C) and light-adapted fluorescence under actinic light (B, D). Kinetics of chlorophyll fluorescence in selected lines are shown in (A, B); image (A) is magnification of the first 50 seconds of image (B). Shown values are averages +/-standard deviations. The quenching responses were quantified by calculating the integrated area under the chlorophyll fluorescence curve in each tested line for dark relaxation (C) and light-adapted fluorescence (D). The genotypes are arranged in order of increasing mean value. For each experiment we used leaf discs from at least 5 independent plants. The experiment was repeated twice with similar results. Statistically significant groups are indicated with letters, which are bolded for lines that have higher remaining chlorophyll fluorescence than wild type. There was a strong correlation between MV-induced quenching of chlorophyll fluorescence and AOX abundance with Pearson correlation values and p-values of 0.7034, 0.0016 for dark-adapted and 0.6518, 0.0046 for light-adapted chlorophyll fluorescence, respectively. The exception was the *AOX1a*-OE line, which was excluded from the calculation.

To gain further insight into the activity of MV at its site of action on the electron-acceptor side of PSI, we performed flash-induced chlorophyll fluorescence imaging on our plant lines (Shapiguzov et al., 2020). The polyphasic rise of chlorophyll fluorescence triggered by a saturating light pulse, also called the OJIP transient, allows attribution of chlorophyll fluorescence to specific domains of the photosynthetic electron transfer chain. The O-J phase is associated with PSII, the J-I phase with the intersystem electron transfer chain and the I-P rise with PSI (Strasser et al., 2004; Stirbet and Govindjee, 2011, 2012; Schreiber and Klughammer, 2021; Schreiber, 2021). As was expected, MV application specifically diminished the I-P rise indicating that the activity of MV caused oxidation of PSI in all the lines (Supplementary Figure 4). However, after an hypoxic atmosphere was introduced by 60-minute flushing of nitrogen gas in darkness, the plant lines demonstrated differences in the extent of the I-P rise (Fig. 5). We calculated φRE1o = 1 − Fi/Fm, which is a parameter sensitive to the described effect of MV on the I-P rise (Strasser et al., 2004; Shapiguzov et al., 2020) for MV-treated samples and corresponding controls. MV still quenched the I-P rise in the wild type, but its impact was significantly diminished in several lines with impaired mitochondria (Fig. 5).

**Figure 5.**
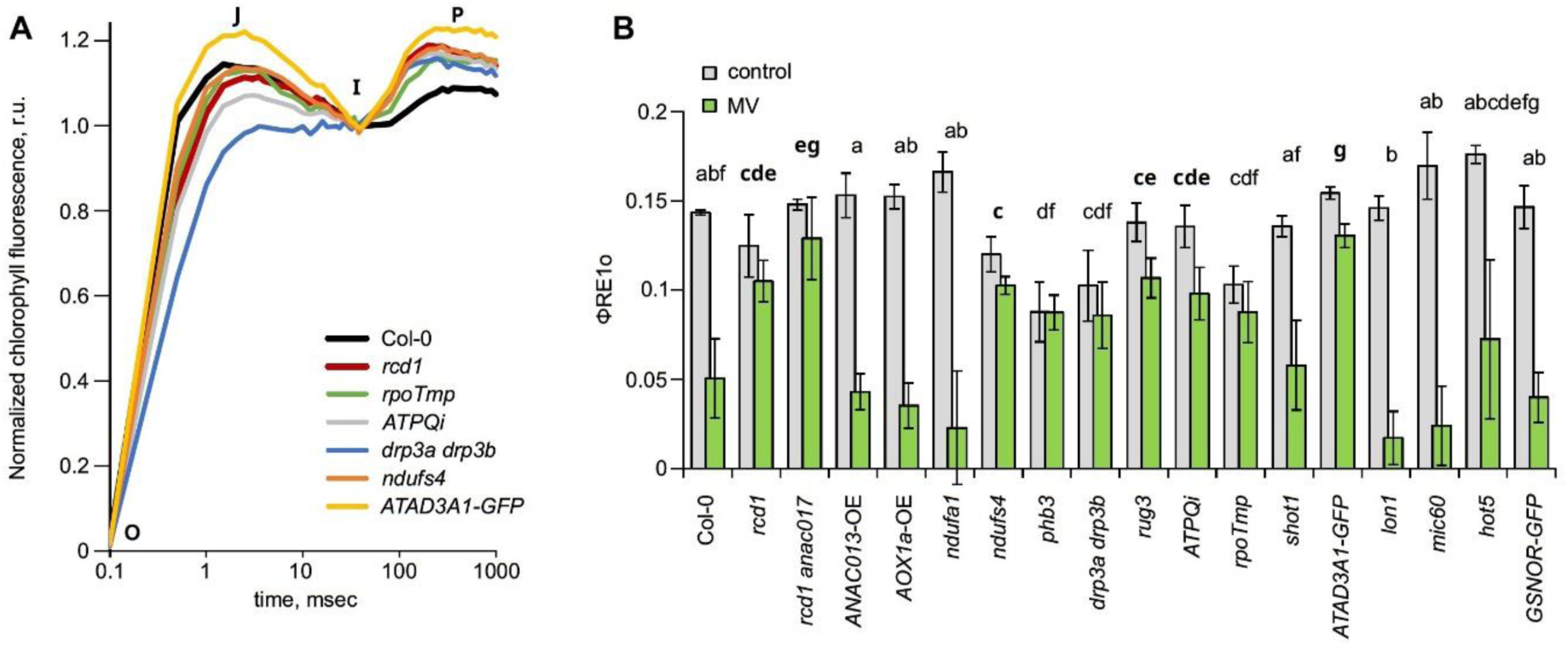
Flash-induced chlorophyll fluorescence in the presence of MV under hypoxic conditions in plant lines with perturbed mitochondrial functions. Flash-induced chlorophyll fluorescence imaging was used for examining the effects of MV directly on the electron-acceptor side of PSI. OJIP kinetics in selected lines under hypoxic conditions in MV-treated leaf discs are shown in (A). Quantification of φRE1o = 1 − Fi/Fm was used as an indicator of the I-P rise, corresponding to PSI activity (B). Shown values are averages +/-standard deviations. Leaf discs from 4 independent plants were assayed. Statistically significant groups are indicated with letters, which are bolded for lines that have higher φRE1o value in the presence of MV than wild type. There was a strong correlation between φRE1o in the presence of MV and AOX abundance, with Pearson correlation value of 0.6385 and p-value 0.006.

Altogether, the above measurements of MV-induced chlorophyll fluorescence quenching demonstrated that the effects of MV on photosynthetic electron transfer and specifically on PSI were modified in the lines with altered mitochondrial functions. However, the differences in MV response only became visible after the plants had been shifted to an hypoxic atmosphere. This suggested that the interaction of altered mitochondria with photosynthetic electron transfer was dependent on availability of O_2_.

### Mitochondrial defects modify consumption and accumulation rates of O_2_ *in vivo*

The observation that the effects of MV on PSI oxidation were modulated by an externally generated hypoxic atmosphere prompted us to study whether altered response of mitochondrial mutants to MV was linked to tissue levels of O_2_. We hypothesized that the changes in respiration caused by mitochondrial defects could affect tissue O_2_ availability, thereby modulating the effects of MV. Thus, we selected a subset of lines with the strongest changes in MV response compared to the wild type and measured their rates of O_2_ consumption in darkness and O_2_ re-accumulation in light. An example of the assay is shown in Figure 6A. We found several lines that had higher O_2_ consumption rates in darkness and lower O_2_ re-accumulation rates in light than in the wild type (Fig. 6B, C). The studies in *rcd1* showed no defects in its chloroplast proteome (Shapiguzov et al., 2019) or O_2_ evolution (Shapiguzov et al., 2020). Thus, the changes in O_2_ re-accumulation rate were likely due to increased reuptake of photosynthetically evolved oxygen not to decreased O_2_ evolution.

**Figure 6.**
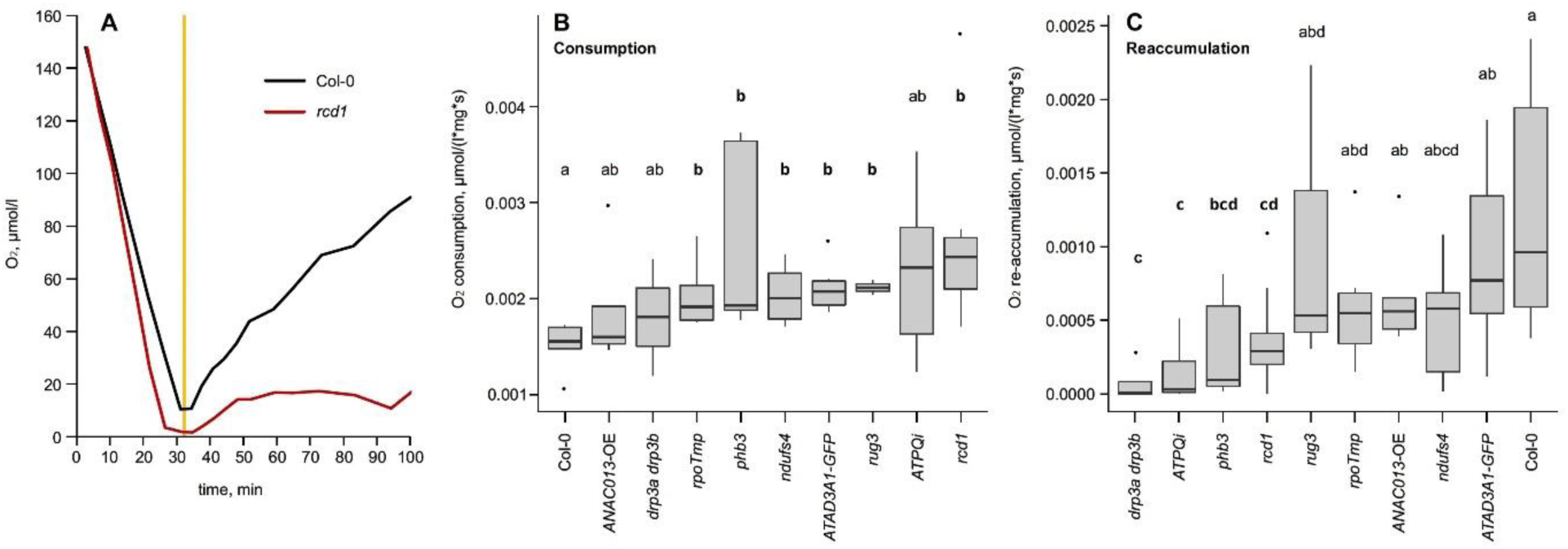
O_2_ consumption and accumulation dynamics in plant lines with perturbed mitochondrial functions. O_2_ dynamics in selected plant lines were examined by measuring O_2_ fluctuations in cuvettes where aerial parts of 3-week-old plate-grown plants were immersed in analysis buffer. An example of this assay performed with Col-0 and *rcd1* is shown in (A), with relative O_2_ consumption rate first recorded in darkness. After O_2_ concentrations approached zero, the light was turned on (yellow vertical line) and O_2_ re-accumulation due to photosynthetic O_2_ evolution was recorded. Consumption (B) and re-accumulation (C) rates of O_2_ were obtained from the kinetics as shown in (A). The genotypes are arranged in order of increasing mean value. Statistically significant groups are indicated with letters, which are bolded for lines that have higher O_2_ consumption or lower O_2_ re-accumulation rates than wild type.

We next performed a similar experiment in the presence of the AOX inhibitor salicylhydroxamic acid (SHAM). SHAM was dissolved in DMSO, thus pure DMSO was added to the untreated controls. Addition of SHAM reduced the above differences between the lines (Supplementary Figure 5). This experiment supported our hypothesis that the differences in O_2_ dynamics were related to different levels of AOX activity.

In the above-described assays we measured exogenous levels of O_2_ that diffused from plant tissues to the outside media in the measurement cuvette. To find out whether we could also observe changed levels of O_2_ inside plant tissues, we probed the O_2_ status inside the central veins of leaf blades *in situ* using oxygen microsensors (Supplementary Figure 6A). Even though no statistical difference was detected in oxygen consumption rates between Col-0 and *rcd1*, oxygen re-accumulation after the start of illumination was suppressed in *rcd1* (Supplementary Figure 6B). This was in line with the measurements of external O_2_ dynamics in seedlings.

The results obtained in all the tested lines demonstrated that AOX abundance strongly correlated with AOX respiration capacity, while all the described MV-related phenotypes correlated with each other (Fig. 7A). When *AOX1a*-OE line was removed from the correlation analyses, a strong positive correlation was observed between AOX abundance, AOX capacity and diverse MV phenotypes (Fig. 7B). Furthermore, resistance to MV-induced photoinhibition showed positive correlation with O_2_ consumption rate in darkness (Fig. 7C). MV-related phenotypes also negatively correlated with O_2_ re-accumulation rate in light, although with low statistical significance. Taken together, these observations suggest a link between AOX activity, tissue O_2_ status and MV-catalyzed Mehler reaction at the electron-acceptor side of PSI (Fig. 7).

**Figure 7.**
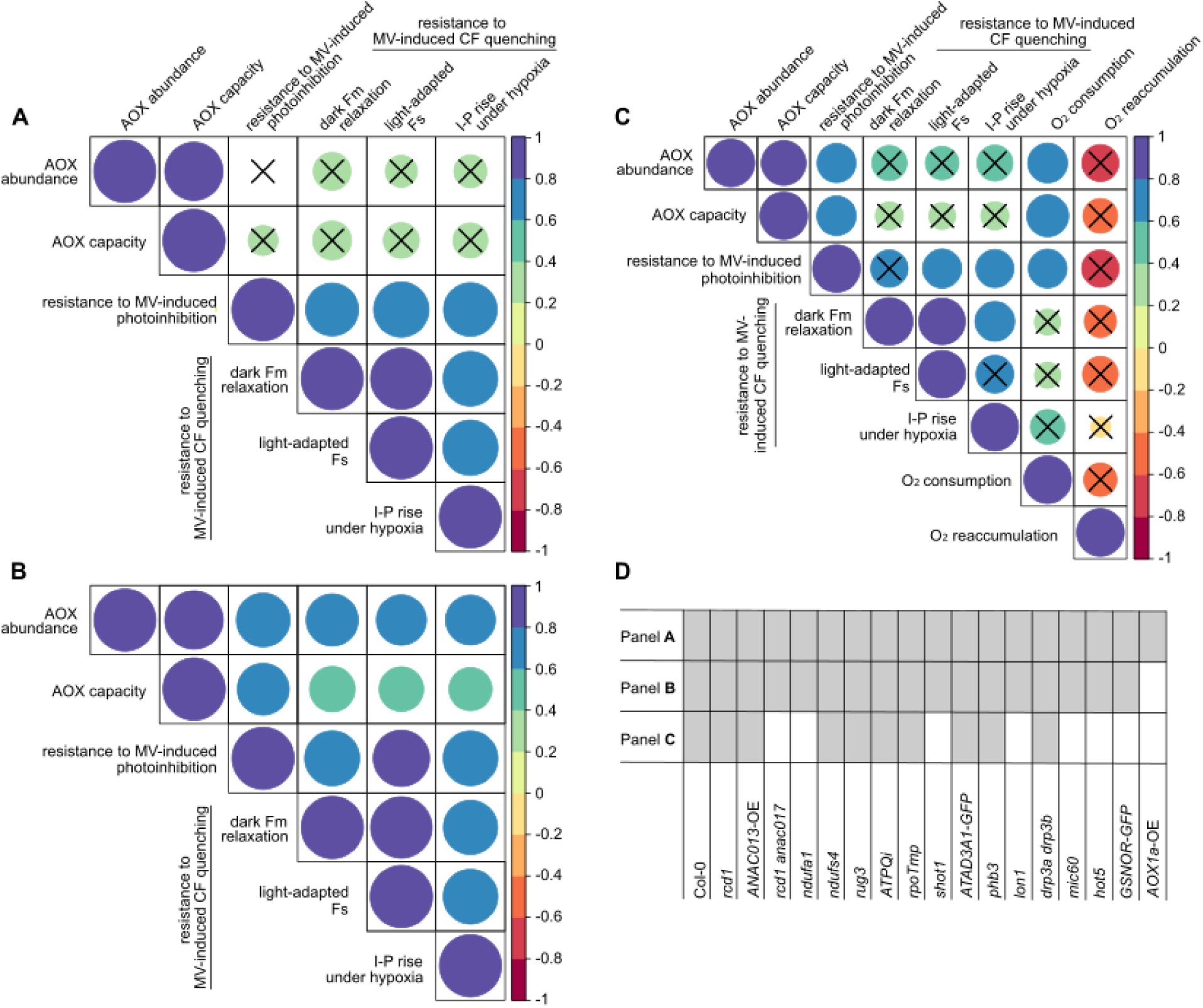
Correlation matrices of chloroplastic and mitochondrial phenotypes in plant lines with perturbed mitochondrial functions. Parameters that were calculated in this study were analyzed using Pearson correlation. Correlations were calculated in tests where all available plant lines were used (A), or with atypically behaving *AOX1a-*OE line excluded (B). A separate matrix was calculated for the selected lines tested in the O_2_ consumption / re-accumulation assay (C). Plant lines included in (A), (B) and (C) are shown in (D). Size and color of the circles indicate strength of correlation, the color scale is shown to the right of each panel. Statistically nonsignificant correlations (p > 0.05) are marked with crosses. For more detail see Supplementary Dataset 1. Raw data and statistics.

## Discussion

### The emerging interaction of plant mitochondria and chloroplasts through intracellular exchange of O_2_

The effects of enhanced mitochondrial respiration and AOX activities on photosynthesis have been intensively addressed. In most studies, the impact of AOXs on photosynthesis was linked to their electron sink activity allowing these enzymes to process reducing equivalents imported to mitochondria from the chloroplast. According to this widely accepted view, AOXs mitigate ROS production both in mitochondria and chloroplasts by reducing redox poise of the ubiquinol pool (Yoshida and Noguchi, 2011; Vanlerberghe, 2013, 2020; Bailleul et al., 2015; Dahal and Vanlerberghe, 2017; Murik et al., 2019; Møller et al., 2021; Oh et al., 2022). O_2_ sink activities of plant mitochondria and AOXs could potentially contribute to diminished cellular ROS production as well. O_2_ sink roles of mitochondria have been discussed in animal cells (Mori et al., 2022). In higher plants, mitochondria and AOXs were implicated in establishment of tissue hypoxia in non-green tissues (Gupta et al., 2009; Rasmusson et al., 2009; Kelliher and Walbot, 2012; Van Dongen and Licausi, 2015). An obvious reason hampering studies in photosynthetically active tissues is the experimental challenges associated with endogenous evolution of photosynthetic O_2_, which masks the effects of mitochondrial O_2_ consumption in light. In this study, we revealed and probed this inter-organelle interaction in special experimental conditions including an hypoxic environment and treatment of plants with MV. MV catalyzes the Mehler reaction, electron transfer from PSI to O_2_. Physiological consequences of MV-catalyzed ROS production at the electron-acceptor side of PSI are relatively well studied (Farrington et al., 1973; Nishiyama et al., 2011; Hawkes, 2014; Shapiguzov et al., 2019). In addition to these more long-term effects (tens-of-minutes to hours range), MV abolishes photosynthetic O_2_ evolution within a few minutes of exposure to light. This is likely caused by suppression of chloroplastic ATP synthase and, as a result, enhanced acidification of the thylakoid lumen and an increase in non-photochemical quenching (Shapiguzov et al., 2020). Importantly, MV does not inhibit CO_2_ evolution, which indicates unaltered mitochondrial respiration. These effects likely explain similarities between MV-induced stress and hypoxia (Shapiguzov et al., 2020). By catalyzing the Mehler reaction, MV quenches chlorophyll fluorescence, thus generating a detectable physiological signal *in planta*. The above physiological features made the use of MV instrumental for this study.

Our results suggest that increased mitochondrial respiration, possibly associated with elevated AOX activity, affects photosynthetic electron transfer and chloroplast functions through diminishing intracellular concentrations of O_2_. O_2_ deficiency affects the Mehler reaction at the electron-acceptor side of PSI. This modulates photosynthetic electron transfer and ROS/redox metabolism of the chloroplast. Further studies are required to address the significance of these findings under physiological conditions, without external hypoxia or MV treatment. Certain phenotypes of the studied lines suggest that this interaction also occurs in natural environments. For example, the *rcd1* mutant is characterized by more reduced redox states of chloroplast thiol enzymes under ambient atmosphere, which is suggestive of the defects in the Mehler reaction (Shapiguzov et al., 2019, 2020).

### Controversial O_2_ sink roles of AOX and COX respiratory pathways

Implication of AOXs in the interaction between mitochondria and the chloroplast Mehler reaction is supported by strong correlations of the corresponding chloroplastic phenotypes with AOX abundance and capacity (Fig. 7), as well as by the observation that application of AOX inhibitor SHAM modifies photosynthetic electron transfer and MV toxicity (Shapiguzov et al., 2019; Pascual et al., 2021).

However, several findings challenge the involvement of AOXs in this organelle interaction. The transgenic *AOX1a*-OE line (Umbach et al., 2005) has KCN-tolerant respiration, indicative of high AOX capacity (Fig. 2; Shapiguzov et al., 2019). However, AOX1a overexpression alone is insufficient to modify wild-type responses of plants to MV (Fig. 7; Shapiguzov et al., 2020). This comes in contrast with the other lines with mitochondrial defects studied here, in which elevated expression of *AOXs*, most likely *AOX1a* among them, correlated with tolerance to MV. One possible reason why overexpression of a single *AOX1a* gene fails to provide MV tolerance versus mutants with constitutive MDS response, is that the latter have a broader transcriptional response with up-regulation of many genes encoding mitochondrial proteins (De Clercq et al., 2013; Shapiguzov et al., 2019). This is in line with the reported functional cooperation between AOX1a and type II NAD(P)H dehydrogenase NDB2 (Sweetman et al., 2019; Sweetman et al., 2022).

Other pieces of evidence that challenge the involvement of AOXs in the studied organelle interaction include the wild-type reactions to MV of the *aox1a*, *aox1c*, and *aox1d* single loss-of-function mutants (Shapiguzov et al., 2020), as well as the indistinguishability of MV-related phenotypes in *rcd1* versus *rcd1 aox1a* double mutant (Shapiguzov et al., 2019; Shapiguzov et al., 2020). These phenomena could be explained by partially redundant functions of five Arabidopsis AOX isoforms, AOX1a, 1b, 1c, 1d, and 2. AOX1a and AOX1d are the most abundant AOXs in leaves and shoots. *AOX1a* and *AOX1d* are also among the most transcriptionally responsive genes in various types of stress (Clifton et al., 2006). Further studies of multiple *aox* loss-of-function mutants are necessary to elucidate O_2_ sink roles of individual AOX isoforms.

Dissecting the exact molecular processes underlying mitochondrial O_2_ sink activities *in vivo* may be challenging due to the high metabolic flexibility of respiration (Sweetlove et al., 2010). One of the factors contributing to functional complexity of mitochondrial O_2_ consumption is metabolism of nitric oxide (NO). Under normoxic conditions AOXs reduce electron leakage from mitochondrial electron transfer chain to nitrite and, hence, diminish NO formation. However, under hypoxia when intracellular environment is over-reduced, mitochondria may enhance NO production, which can inhibit the COX pathway. Interestingly, however, increased NO emission under hypoxia was observed in *AOX* overexpressor lines in tobacco (Jayawardhane et al., 2020) and Arabidopsis (Vishwakarma et.al., 2018), suggesting the existence of a complicated interplay of AOX and COX respiratory pathways through NO metabolism. We used *hot5* mutant and *GSNOR-GFP* line with unbalanced NO homeostasis (Lee et al., 2008). In our assays, these lines were indistinguishable from the wild type, suggesting that NO was not playing a major role in the studied organelle interaction.

Extensive metabolic analyses in *rcd1* revealed elevated respiratory metabolic flux (Shapiguzov et al., 2019) and changed levels of numerous primary metabolites many of which may act as respiratory substrates (Sipari et al., 2020). Thus, involvement of specific AOX isoforms and/or COX respiratory pathway in the studied interaction may be flexibly interchangeable in different physiological contexts.

### Possible implications and open questions

Our observations open new research possibilities. The exact molecular processes contributing to mitochondrial O_2_ sink effects are yet to be determined. The effects of enhanced O_2_ sinks, including elevated mitochondrial respiration, on photosynthetic electron transfer and redox states of thiol redox enzymes need to be addressed, with particular interest in chloroplast thioredoxin networks that are known to depend on Mehler reaction as the electron sink. Further studies will address the effects of mitochondrial O_2_ consumption and other O_2_ sinks on ROS signaling pathways in various cellular compartments including mitochondria and chloroplasts, but also other cellular compartments including apoplast and nucleus (Shapiguzov et al., 2012; Waszczak et al., 2018). The implication of the described interaction in stress reactions, in particular those related to hypoxia need to be addressed. For example, mitochondrial MDS signaling pathway merges on the one hand with chloroplast ROS signaling at the level of the nuclear protein RCD1 (Shapiguzov et al., 2019). On the other hand, MDS pathway is linked to transcriptional reprogramming induced by hypoxia (Shapiguzov et al., 2020). The role of the described O_2_ sink effects in these pathways need to be evaluated. Additionally, our study opens new possibilities in phenotyping plant respiration. Studying plant mitochondrial respiration *in situ* is challenging because of the absence of detectable spectroscopic signatures. Here we present evidence that in certain physiological situations mitochondrial respiration impacts chlorophyll fluorescence. This provides a technical possibility to deduce the traits related to enhanced plant respiration from chlorophyll fluorescence analyses. This can serve as a basis to develop future tools for imaging plant respiration.

## Materials and methods

### Plant material and growth conditions

*Arabidopsis thaliana* accession Columbia-0 (Col-0) was used in all experiments. The plants were grown on either 1:1 mix of soil and vermiculite in 12 h photoperiod with 200-250 μmol m^−2^ s^−1^ illumination (low light growth conditions) or on MS plates (MS-medium, gel, MES hydrate) in 12 h photoperiod with (growth chamber condition) illumination. The following mutants were used in the experiments: knockout mutants *rcd1-4* (AT1G32230), *rcd1-1 anac017* (AT1G32230/AT1G34190; Ng et al., 2013; Shapiguzov et al., 2019), *shot1-2* (AT3G60400; Kim et al., 2012), *hot5-2* (AT5G43940; Feechan et al., 2005; Lee at al, 2008), *rpoTmp-1* (AT5G15700; Kühn et al., 2009), *lon1-2* (AT5G26860; Rigas et al., 2009), *drp3a drp3b* (AT4G33650/AT2G14120; Fujimoto et al., 2009), *ndufa1* (AT3G08610; Meyer et al., 2011), *ndufs4* (AT5G67590; Meyer et al., 2009), *rug3-1* (AT5G60870; Kühn et al., 2011), *phb3* (AT5G40770; Van Aken et al., 2007), *mic60-1* (AT4G39690; Michaud et al., 2016), as well as overexpression lines *atad3a1 atad3b1*: *ATAD3A1-GFP* (AT3G03060; Kim et al., 2021), *GSNOR-GFP* (AT5G43940; Feechan et al., 2005; Lee at al, 2008), *AOX1a* overexpressor (AT3G22370; Umbach et al., 2005), *ANAC013* overexpressor (AT1G32870; De Clercq et al., 2013), and RNAi mutant ATPQi (AT3G52300; Liu et al., 2021). All mutants and lines are in Col-0 background.

### Protein abundance measurements

Total AOX abundance was determined by immunoblotting. Plants were grown on MS plates for 18 d, collected whole, ground in liquid nitrogen, and proteins were extracted by incubating the samples for 20 min at 37 °C in lysis buffer (2% SDS, 20 mM Tris-HCl pH 7.8 supplemented with protease inhibitor cocktail P9599, Sigma-Aldrich). After centrifugation for 5 min at 15 000 g, the samples were suspended in Laemmli buffer. Protein amounts were normalized to fresh plant mass. The extracts were separated on a 12% SDS-PAGE gel and transferred to Bio-Rad Immun-blot PVDF membrane. AOX amounts were detected with an AOX antibody (Anti-AOX1/2 antibody AS04 054, dil. 1:5000, Agrisera) together with an HRP-tagged secondary antibody (ECL anti-rabbit IgG LNA934V/AH, GE Healthcare), and detected using a UVP Biospectrum 610 imaging system. For total protein estimation, the membrane was stained with amido black. Band intensities were calculated with the gel analyzer in ImageJ. Quantification was done using 4 independent immunoblots.

### Measurements of O_2_ consumption rates in presence of respiration inhibitors

For measurements of plant O_2_ consumption in the presence of respiration inhibitors, plants were grown for 18 d on MS plates, arial parts were cut off, weighed (approximately 20 mg/sample), and placed in an Oxygraph measurement chamber (Hansatech instruments) in air-saturated liquid MS medium. Oxygen consumption in the dark was monitored with a Clark-type oxygen electrode until it stabilized, for approximately 15 min. After this, changes in oxygen consumption were measured in the presence of respiration inhibitors by injecting potassium cyanide (KCN, 4 mM) into the measurement chamber, waiting for the signal to stabilize, then adding SHAM (2 mM) in the same manner. Respiratory parameters were processed with the Oxygraph plus software, and relative oxygen consumption rates were determined by normalizing the rates to plant mass. Six biological replicates were analyzed.

### Chlorophyll fluorescence imaging

MV-induced photoinhibition and quenching of Fm and light-adapted chlorophyll fluorescence was assessed by chlorophyll fluorescence imaging using IMAGING-PAM M-Series (Heinz Walz, Effeltrich, Germany). Photoinhibition protocols are described in (Shapiguzov, Kangasjärvi, 2022). In brief, leaf discs were floated overnight in darkness on solutions with or without MV, to facilitate uptake of the chemical. After incubation, MV toxicity was induced by repetitive 1-hour light periods (450 nm, 80 µmol m^−2^ s^−1^) each followed by a 20 min dark period, then Fo and Fm measurement. The resulting decay of maximal quantum yield of Photosystem II (PSII) was calculated as Fv/Fm = (Fm-Fo)/Fm. For each experiment we used leaf discs from at least 4 independent plants. The experiment was repeated twice with similar results. Chlorophyll fluorescence quenching routines are described in (Shapiguzov et al., 2020). In brief, leaf discs were prepared as above. An hypoxic environment was introduced by flushing nitrogen gas over the leaf discs for 20 min in darkness. Imaging of chlorophyll fluorescence was conducted under a saturating light pulse (Fm relaxation measurement) followed by low-intensity actinic light (450 nm, 80 µmol m^−2^ s^−1^; Fs measurement). For each experiment we used leaf discs from at least 5 independent plants (for MV experiments) or 3 independent plants (for control experiments). The Fm relaxation experiment was repeated three times with similar results, and the Fs experiment was repeated twice with similar results. For flash-induced chlorophyll fluorescence imaging we used PlantScreen SC Mobile System equipped with ultra-fast CMOS camera TOMI 3 with image acquisition of maximal frame rate 20 µs (Photon Systems Instruments, Drásov, Czechia). Imaging protocols are described in (Shapiguzov et al., 2020). In brief, plant leaves or leaf discs were placed in ultrapure water with added 0.05% Tween-20, in the presence or absence of 2 µM MV. Leaves were incubated in darkness to uptake MV for approximately 16 h before imaging. To minimize the effects of MV on photosynthesis, precautions were taken not to expose plant material to any light prior to measurements. For anoxic measurements, samples were placed in an airtight container that was continuously flooded with nitrogen gas and the measurements were performed through the transparent cover.

### Shoot tissue O_2_ consumption and production

Shoot tissue oxygen consumption was measured using the MicroRespiration system (Unisense, Denmark). Plants were grown on MS plates for approximately 3 weeks. Their aerial parts were cut off, weighed to obtain fresh mass, and inserted into 2 ml glass vials, which were filled with analysis buffer (Smart and Barko solution, 0.6 mM MgSO_4_, 0.8 mM CaCl, 1 mM KHCO_3_ supplemented with 5 mM MES monohydrate). The analysis buffer was partially deoxygenated by mixing at 1:1 ratio O_2_-saturated buffer and buffer that had been deoxygenated by bubbling it with N_2_ gas, resulting in an initial O_2_ concentration of approximately 150-200 µmol/l. The vials were placed in a rack inside a constant temperature bath (37 °C) and O_2_ concentrations were monitored through a capillary in the glass lid with an optode (OPTO-MR, Unisense Denmark) while the plants were treated to the following regime: vials were kept in the dark until O_2_ concentrations in all vials had been reduced to <10 µmol/l, or until the O_2_ concentration in at least one vial had remained at <1 µmol/l for at most 20 min. After this, the vials were illuminated (30 μmol photons m^−2^ s^−1^) until the O_2_ concentrations within vials had plateaued, or at most 2 h. O_2_ concentrations were recorded with RATE (Unisense, Denmark), and O_2_ consumption in darkness as well as O_2_ production in light were normalized to fresh sample mass. At least 5 biological replicates were used, with the exception of *drp3a drp3b* (3 replicates) and *rug3* (3 replicates). For measurements in the presence of SHAM, either DMSO (control) or 8 mM SHAM (treatment) was added to the analysis buffer at the beginning of the measurement. Contamination between samples was avoided by rinsing the O_2_ optode in fresh buffer after SHAM-containing samples were measured.

### Leaf tissue O_2_ dynamics

O_2_ status of intact leaves submerged in deoxygenated medium was measured using Clark-type O_2_ microsensors. Whole leaves were excised, kept submerged in 0.05% Tween-20 solution overnight (12-16 h) in the dark, and placed over a firm foam with the main vein facing upwards. A solid piece of plastic with a hole in the center was positioned over the leaf, the sandwiched leaf was attached on a metal mesh firmly by using rubber bands and placed inside an aquarium (200×100×100 mm, in total 2 L). An O_2_ microsensor (OX-10, Unisense, Denmark) was inserted 80-100 µm into the main vein of the leaf lamina, with 0 indicating the tissue surface. Stagnant, deoxygenated buffer solution (0.1% agar) was added to the aquarium and petiole O_2_ concentration was recorded using LOGGER (Unisense, Denmark) at a rate of 1 hz. The initial part of the measurement was conducted in darkness, until the O_2_ concentration declined and remained at a steady state below 5 µM. Then, the light was turned on (ca. 15 µmol photons m^-2^ s^-1^) and O_2_ concentration was measured until it reached a plateau. The O_2_ concentration and temperature of the external medium were monitored using an O_2_ minioptode (OPTO-MR, Unisense, Denmark) and a temperature probe (ZNTC, Unisense, Denmark). The positioning of the microsensor inside the petiole was aided by a motorized micromanipulator controlled by LOGGER and visually aided with a dissection microscope (WILD M3B, Leica, Switzerland).

### Statistical analyses

Statistical analyses of all results were conducted with MVApp (Julkowska et al. 2017). Differences between variances were estimated with ANOVA or Kruskal-Wallis, depending on the normality and equality distributions of the samples. Correlation was measured with Pearson. Standard cut-off limit for statistically significant p-value (0.05) was used for all experiments. The box and whisker plots included in figures show the median (horizontal line), the first and third quartiles (box), minimum and maximum (whiskers), and outliers (dots). Raw data and statistics are presented in Supplementary Dataset 1.

## Author contributions

A.S., M.P., O.P., E.V. and L.L.P.O. conceived and designed experiments. M.P., L.L.P.O., M.K. and A.S. carried out experiments. M.P., O.B., L.L.P.O., M.K., K.F., E.V., O.P. and A.S. analyzed the results. A.S. and M.P. wrote the article. All authors read and contributed to the final article.

## Funding

This work was supported by the Centre of Excellence in Tree Biology, Research Council of Finland (decision 346140; Alexey Shapiguzov).

## Acknowledgements

The authors dedicate this study to the memory of Prof. Jaakko Kangasjärvi. We thank Prof. Romy Schmidt-Schippers for the advice on the studies of hypoxia and for her critical comments on the manuscript. We thank Dr. Mikael Brosché and Dr. Julia Vainonen for their help in preparation of this manuscript.

## Supplementary Figures

**Supplementary Figure 1.**
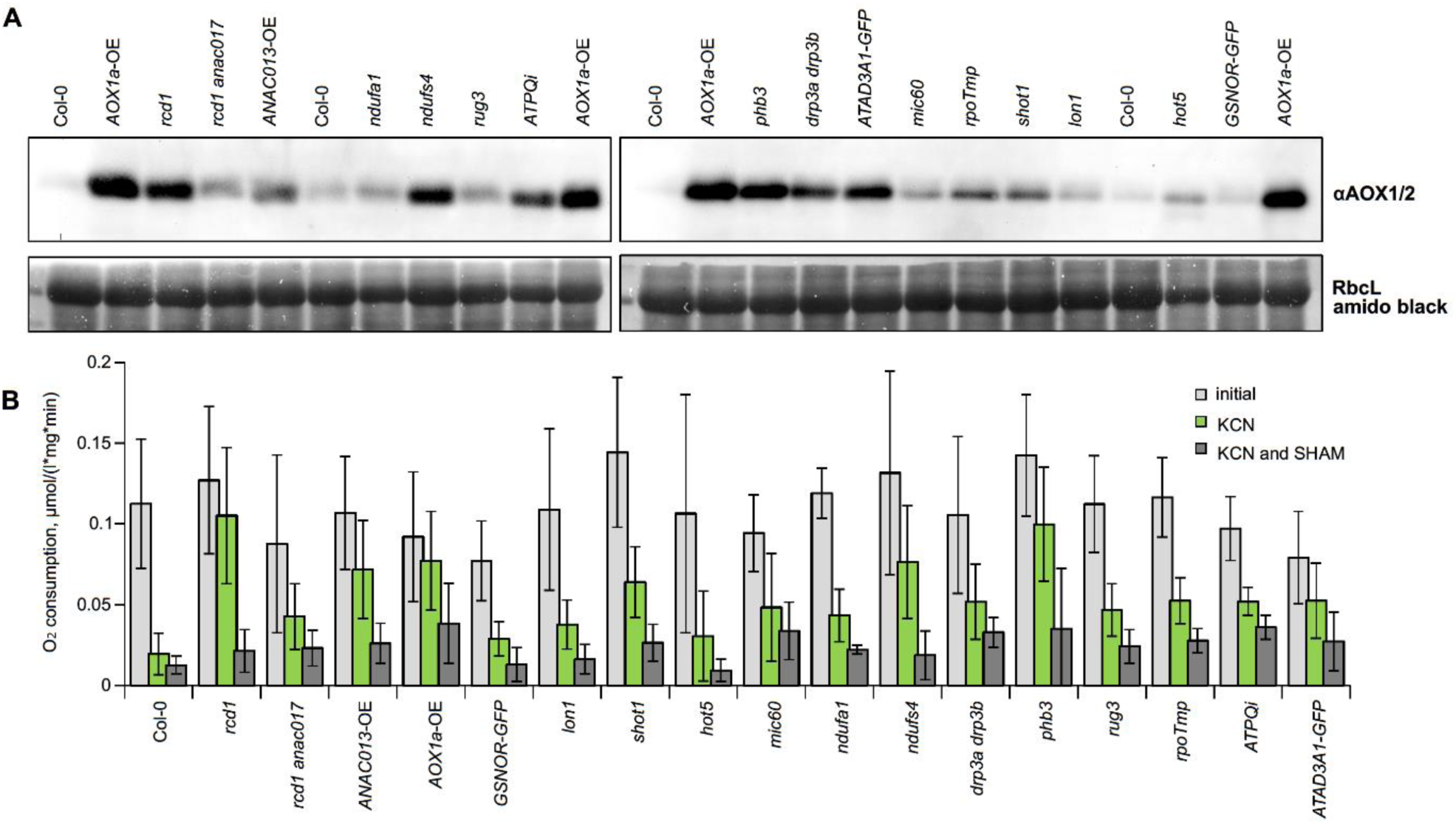
AOX1 abundance and capacity measurements. Representative anti-AOX1 immunoblot and corresponding total protein abundance (A). Inhibition of O_2_ consumption after addition of inhibitors KCN and SHAM (B), shown values averages +/-standard deviations.

**Supplementary Figure 2.**
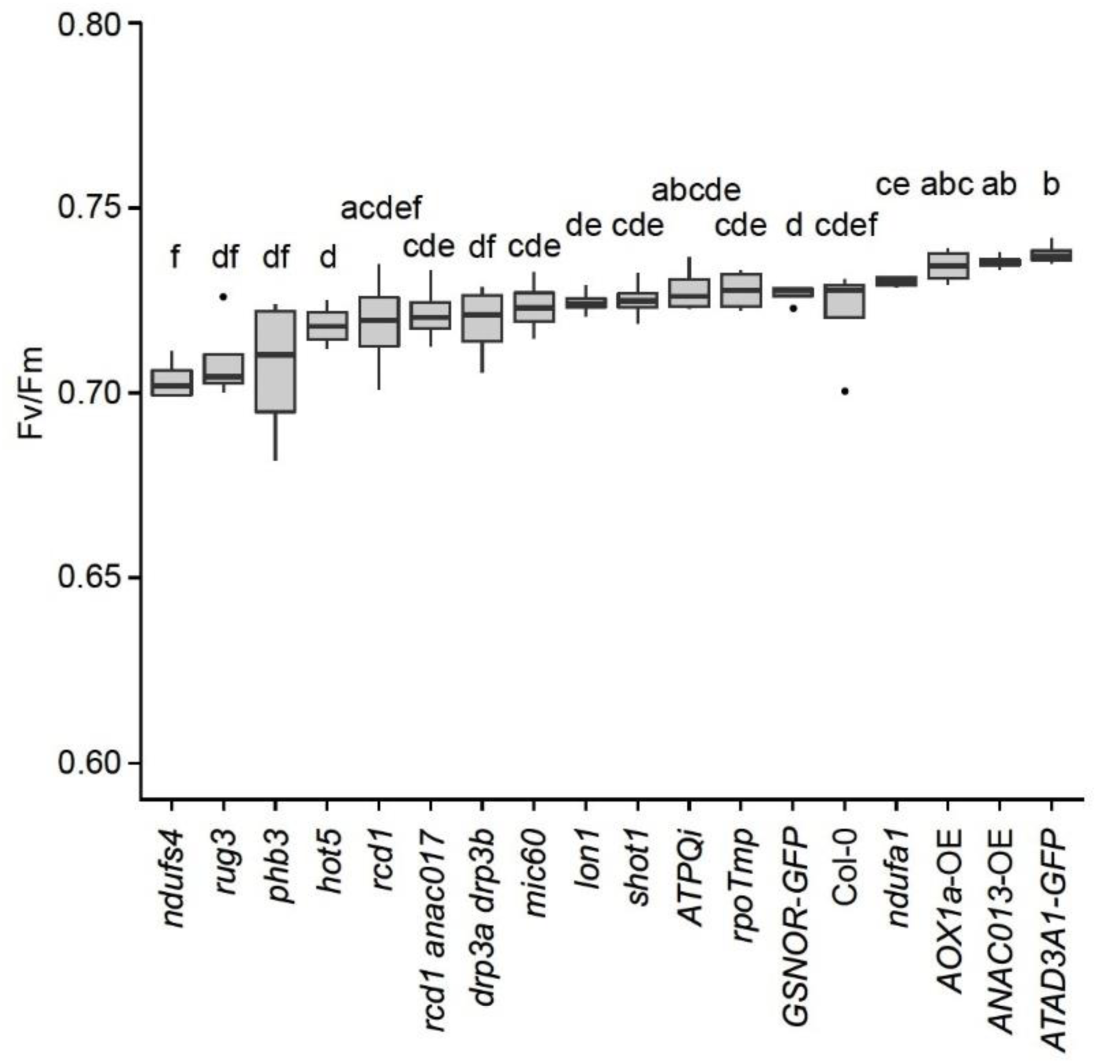
Maximal PSII quantum yield of MV-treated plants prior to light exposure. The genotypes are arranged in order of increasing mean value. Statistically significant groups are indicated with letters.

**Supplementary Figure 3.**
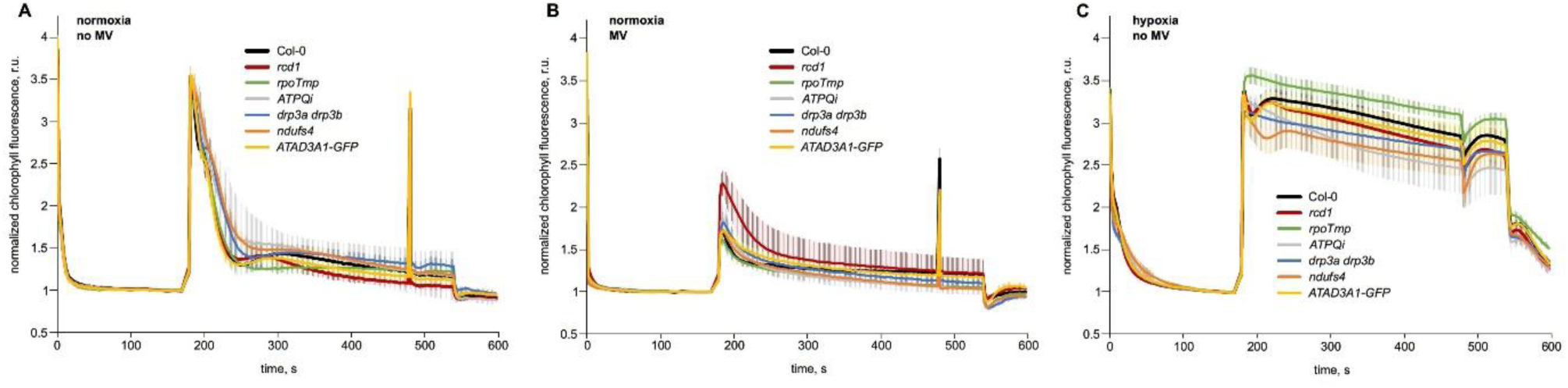
Quenching of chlorophyll fluorescence under control conditions. In addition to the measurements reported in Fig. 4, chlorophyll fluorescence was measured in normoxic conditions in the absence (A) or presence (B) of MV, and under hypoxia without MV (C).

**Supplementary Figure 4.**
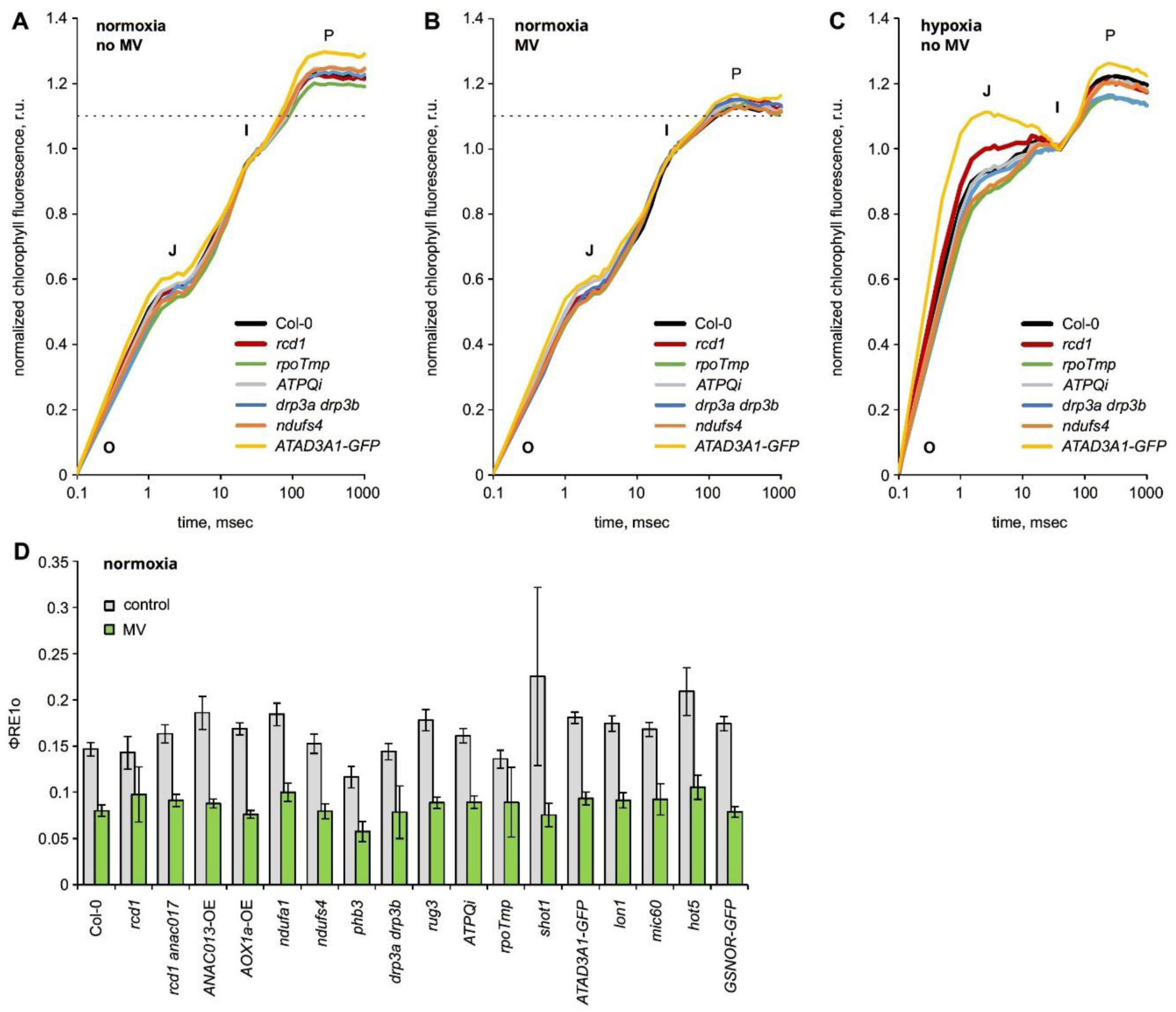
Flash-induced chlorophyll fluorescence measurements in control conditions. In addition to the conditions used in Fig. 5, OJIP kinetics of chlorophyll fluorescence was measured under normoxic conditions in the absence (A) or presence (B) of MV, as well as under hypoxia without MV (C). φRE1o = 1 − Fi/Fm was determined for normoxic conditions (D).

**Supplementary Figure 5.**
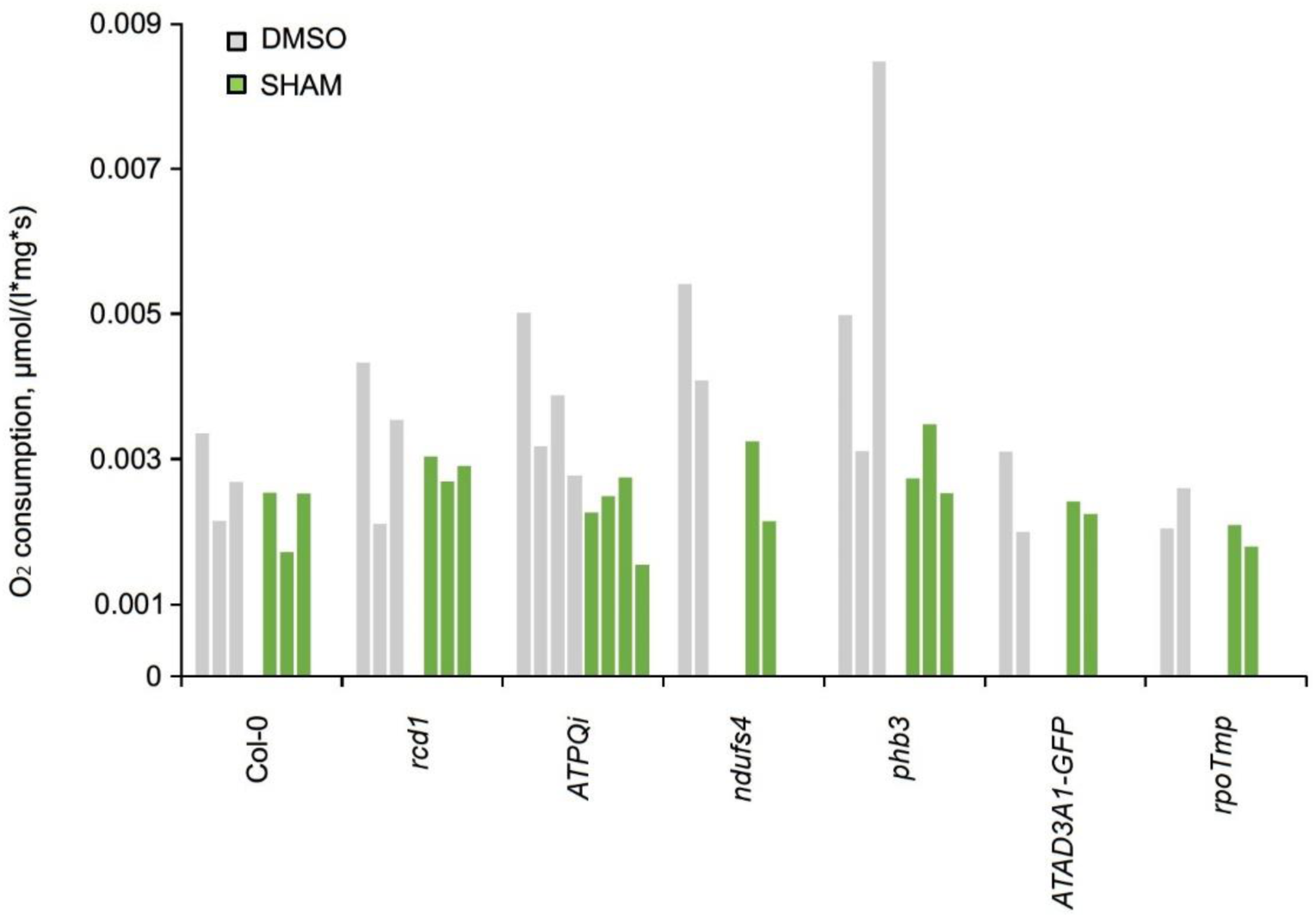
O_2_ consumption dynamics in plant lines with perturbed mitochondrial functions in the presence of AOX inhibitor SHAM. Oxygen consumption rates were determined similarly to Fig. 6 in the presence of DMSO (control) and SHAM. Each bar represents an individual measurement.

**Supplementary Figure 6.**
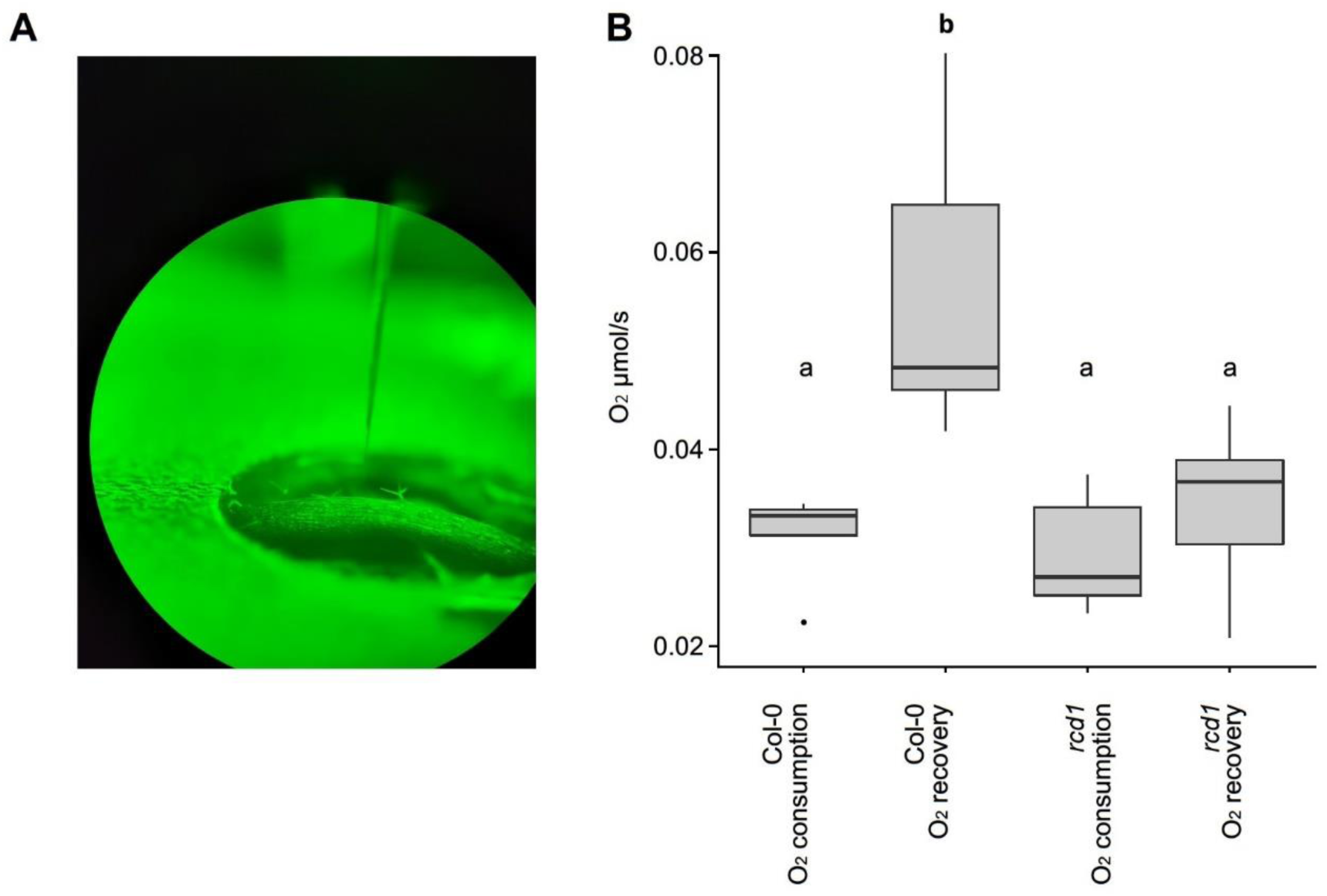
Dynamics of O_2_ measured inside leaf tissues with microsensors. Leaves of Col-0 and *rcd1* plants were inserted into deoxygenated medium and probed with microsensors. A microscope image of the microsensor being inserted into the leaf is shown in (A). Rates for O_2_ consumption (in darkness) and recovery (in light) were measured from inside the central vein of the leaf (B). Statistically significant groups are indicated with letters.

